# The sponge *Hymeniacidon perlevis* is introduced and widespread in the Western Atlantic

**DOI:** 10.64898/2026.08.03.741563

**Authors:** Thomas L. Turner, Humberto F. M. Fortunato, Gisele Lôbo-Hajdu, Juliet M. Wong

## Abstract

The marine sponge *Hymeniacidon perlevis* is a globally-distributed species, likely because it has been extensively spread through human means. The status of this species in the Western Atlantic has been difficult to discern due to the presence of a morphologically similar species *Hymeniacidon heliophila.* Here, we have collected *Hymeniacidon* specimens from the *H. heliophila* type location and compared their morphology and genotypes to material from Europe, North and South America, and publicly available datasets. Specimens from the type location closely matched the original description of *H. heliophila*, and were morphologically indistinguishable from *H. perlevis*. Multilocus sequencing data confirm that these samples are indistinguishable from *H. perlevis*, and that the name *H. heliophila* should therefore be considered a junior synonym of *H. perlevis*. Analysis of a global cox1 dataset further revealed significantly higher genetic diversity in Europe than in the remainder of the world combined, supporting a European origin for the species and suggesting that all other genotyped populations are introductions. In the Western Atlantic, we confirm the presence of *H. perlevis* in the Gulf of Mexico, but contrary to some past reports, are unable to confirm its presence in the Caribbean. Our results clarify the taxonomy, biogeography, and recent history of one of the world’s most widespread marine sponges.

## Introduction

The marine sponge *Hymeniacidon perlevis* is a globally-distributed species. It was first described from southwest England over 200 years ago (Montagu 1814), and it remains common in the intertidal and shallow waters of Europe (Erpenbeck and van Soest 2002; Mahaut et al. 2013; Regueiras et al. 2019). Recently, genetic data have confirmed that the species has spread far beyond Europe, with robust populations in a remarkable number of locations. In the Atlantic, sequencing confirms its presence from the US states of Florida (Erpenbeck et al. 2007) and Alabama (Erwin et al. 2011), Argentina (Gastaldi et al. 2018), and South Africa (Samaai et al. 2022). In the Pacific, it is known from Korea (Park et al. 2007), China (Jun et al. 2015), Japan (Hoshino et al. 2008), the US state of California (Fuller and Hughey 2013), and the Canadian province of British Columbia (Harbo et al. 2021). Morphological data also suggest its presence in additional regions, including Italy (Mercurio et al. 2023), New Zealand (Bergquist 1961), and Peru (Cóndor-Luján et al. 2023), though the morphological similarity of other species of *Hymeniacidon* makes genetic confirmation of these populations desirable.

Several lines of evidence indicate that this remarkable distribution is the result of human-mediated introductions in many of these locations. First, the species has lecithotrophic larvae that generally settle within a few days, making long-distance dispersal unlikely (Xue et al. 2009). Second, populations have been documented expanding their ranges over historical timescales (Fuller and Hughey 2013; Harbo et al. 2021). Consistent with a hypothesis of human introduction, *H. perlevis* is often found growing on human-made structures in highly modified habitats such as floating docks in harbors, where a large fraction of the marine invertebrate community is non-native (Erwin et al. 2011; Mercurio et al. 2023; Turner et al. 2025). However, the species occupies a broad ecological niche and also occurs in relatively pristine habitats, including California’s kelp forests where few other introduced invertebrates are known to have established (Turner 2020; Turner et al. 2025). It is also commonly found in wave-swept rocky intertidal habitats (Samaai et al. 2022; Turner et al. 2025) and on oysters and other hard substrates in bays and mud flats (Fuller and Hughey 2013; Harbo et al. 2021). In addition to its ecological breadth, *H. perlevis* also spans a broad latitudinal range, in both Atlantic and Pacific oceans, occurring in both cold-temperate and warm-temperate regions (Turner 2020; Turner et al. 2025). However, its status in tropical waters remains unclear. Ecological niche modeling suggests that sea surface temperature may be the primary factor limiting its spread (Samaai et al. 2022), but this inference is only as reliable as the occurrence data used to build the model and could change if tropical populations are genetically confirmed.

One region where the status and distribution of *H. perlevis* remains highly uncertain is the Western Atlantic. Until recently, the species was not thought to occur there (Muricy et al. 2011; Fortunato et al. 2020), and regional sponges matching its morphology were instead assigned to *H. heliophila* (Wilson 1911). *H. heliophila* was described by H. V. Wilson in 1911 from specimens collected at Pivers Island in Beaufort, North Carolina, on the Atlantic coast of the United States (Wilson 1911). Subsequent authors applied the *H. heliophila* species concept to *Hymeniacidon* populations across a broad geographic range in the Western Atlantic. As a result, the Global Biodiversity Information Facility (gbif.org) lists vouchered samples of *H. heliophila* from New England in the Northwest Atlantic to Argentina in the Southwest Atlantic. The species has also been reported as widespread throughout the Caribbean (Diaz et al. 1993), with published occurrences that include the Guyana Shelf (van Soest 2017), and Curacao (Diaz et al. 1993), as well as reports from the Gulf of Mexico (Erwin et al. 2011; de la Cruz-Francisco 2025).

A recent compilation of genetic data demonstrated that many specimens previously identified as *H. heliophila* are actually *H. perlevis* (Turner 2020). Samples from Alabama in the Gulf of Mexico and from the Atlantic coasts of Florida and Argentina were genetically indistinguishable from *H. perlevis* (Gastaldi et al. 2018; Turner 2020). One specimen previously identified as *H. heliophila* from North Carolina was likewise genetically indistinguishable from *H. perlevis*, although it was collected from a different part of the state than the *H. heliophila* type locality (Turner 2020). These findings raise the possibility that the sponges examined by Wilson in Beaufort Harbor were themselves an introduced population of *H. perlevis*. If so, the name *H. heliophila* would not be valid and should be treated as a junior synonym of *H. perlevis*. The ecological and morphological observations presented in Wilson’s original description are consistent with this interpretation; nothing in that description distinguishes *H. heliophila* from what is now known about the morphology and ecology of *H. perlevis*. This would not be the first case of a nominal species being synonymized with *H. perlevis*: the World Porifera Database currently lists 19 additional names that are considered junior synonyms of this widespread and morphologically variable species (de Voogd et al. 2026). Determining whether the name *H. heliophila* is valid is a crucial step toward understanding both the introduction history of *H. perlevis* in the Western Atlantic and the ecological breadth of the species globally.

Though previous work has confirmed that *H. perlevis* is present in the Western Atlantic, these studies stopped short of formally recommending that *H. heliophila* be treated as a synonym of *H. perlevis* because several uncertainties remained. First, no genetic data were available from the type locality. Wilson did not preserve the specimens he examined, and no type material exists. We are also unaware of any historical vouchers from Beaufort Harbor. Second, some specimens previously identified as *H. heliophila* are genetically distinct from *H. perlevis*, indicating that at least one additional morphologically similar *Hymeniacidon* species occurs in the region. These outlying samples were collected from Key Largo, in Florida, and identified by Belinda Glasby as *H. heliophila* (Turner 2020). Sequence of the 18S and 28S nuclear ribosomal loci from these samples was highly divergent from all *H. perlevis* sequences, and is clearly from a different species (Turner 2020).

To better understand the introduction history of *H. perlevis* and the diversity and biogeography of Western Atlantic sponges, we collected new *Hymeniacidon* specimens from Beaufort Harbor and present genetic and morphological data from those samples here. By combining these data with newly collected material from Brazil, California, and other locations, we provide a clearer picture of the distribution and eco-evolutionary history of one of the world’s most widespread marine sponges.

## Methods

### Samples and collections

Samples of *Hymeniacidon* were collected from two locations in the Beaufort Harbor estuary, North Carolina, USA: the Duke Aquafarm (n=4 from floating oyster cages, n=8 from oysters on intertidal mudflats immediately adjacent to the cages) and the Rachel Carson Reserve (from oysters on intertidal mudflats, n=12). Additional samples were investigated from a variety of locations including Rio de Janeiro, Brazil; Virginia, USA; California, USA; Puerto Rico; Belize; Barbados; and the Galapagos Islands, Ecuador. California samples include both newly collected material and samples reported in past work (Turner et al. 2025); other samples were acquired on loan from the National Museum of Rio de Janeiro (voucher numbers with MNRJ) and the Smithsonian Museum (voucher numbers with USNM). The full list of samples examined, with associated metadata including GenBank accession numbers and collection information, is given in Table S1.

### Morphology

Spicules were examined after digesting sponge subsamples in bleach or nitric acid. Skeletal architecture was characterized by hand cutting tissue sections and digesting them in a mixture of 97% Nuclei Lysis Solution (Promega; from the Wizard Genomic DNA Purification kit) and 3% 20mg/ml Proteinase K (Promega). This digestion eliminates cellular material while leaving the spongin network and spicules intact. Tangential sections of the ectosome were made directly in fixed specimens. Spicule measurements were made on digital images using ImageJ (Schneider et al. 2012) after calculating the number of pixels per mm with a calibration slide.

Spicule length was measured as the longest possible straight line from tip to tip, even when spicules were curved or bent. Spicule width was measured at the widest point, excluding swollen tyles. All spicule measurements are available as raw data in Table S2. Also included are measurements that were made by others and previously reported only as summary statistics, but generously shared by the authors of previous works via personal communication (Harbo et al. 2021; de la Cruz-Francisco 2025). Visual inspection and previous comparisons of *Hymeniacidon* species in California (Turner et al. 2025) suggested that the variability in spicule length and width may be a useful trait for differentiating *Hymeniacidon* species in the region. Spicule size variability was quantified for each sample as the root mean square (RMS) Euclidean distance of individual spicules from the specimen centroid in two-dimensional length–width space, where the centroid was defined by the mean spicule length and mean spicule width of that sample:

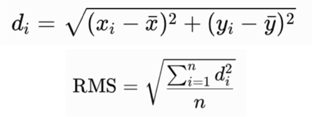

Where x and y are length and width, and RMS is the root mean square dispersion.

### Genotyping

The cox1 locus was sequenced at the Folmer barcoding region using the following primers: (LCO1490: 5’-GGT CAA CAA ATC ATA AAG AYA TYG G-3’; HCO2198: 5’-TAA ACT TCA GGG TGA CCA AAR AAY CA-3’) (Folmer et al. 1994). Three primer sets were used to amplify regions of the nuclear ribosomal locus: the ∼800 bp D1–D2 region was amplified using primers Por28S-15F (5’- GCG AGA TCA CCY GCT GAA T-3’) and Por28S-878R (5’-CAC TCC TTG GTC CGT GTT TC-3’); the ∼800 bp D3–D5 region was amplified using primers Por28S-830F (5’-CAT CCG ACC CGT CTT GAA-3’) and Por28S-1520R (5’-GCTAGT TGA TTC GGC AGG TG-3’) (Morrow et al. 2012). The ITS region was amplified using primers ITS-F (5’-TCA TTT AGA GGA AGT AAA AGT CG-3’) and ITS-R (5’-GTT AGT TTC TYT TCC TCC GCT T-3’) (Lôbo-Hajdu et al. 2004).

PCR at the cox1 locus was performed using the following conditions: 95°C for 3 min, followed by 35 cycles of 94°C for 30 sec, 52°C for 30 sec, 72°C for 60 seconds, followed by 72°C for 5 minutes; for the 28S amplicons the annealing temperature was 52°C; for the ITS region, conditions were changed to 56°C annealing temperature and only a 45 sec extension time. PCR on Brazilian samples was performed in a total volume of 30 μL containing: 0.75 U of GoTaq DNA polymerase (Promega), 1.5 mM MgCl_2_, 0.133 mM dNTPs (Promega), 0.16 μM of each primer, and approximately 20–100 ng of template DNA. PCR on all other samples was performed in 50 μl reactions using the following recipe: 24 μl nuclease-free water, 10 μl 5x PCR buffer (Gotaq flexi, Promega), 8 μl 7.5mM MgCl_2_, 1 μl 10 mM dNTPs (Promega), 2.5 μl of each primer at 10 μM, 0.75 bovine serum albumin (10 mg/ml, final conc 0.15 mg/ml), 0.25 μl GoTaq polymerase, 1 μl template. Blastn was used to verify that the resulting traces were of sponge origin. All sequences have been deposited in GenBank; accession numbers are shown in Table S1.

### Phylogenies

We used blast to assemble additional sequences of *H. perlevis*, other *Hymeniacidon* species, and assorted outgroups. Samples from the barcoding of life database (BOLD) that were annotated as *H. perlevis* or *H. heliophila* were also included if they were not also available from GenBank. Sequences were aligned using MAFFT v.7 (Katoh et al. 2019), and phylogenies were estimated with maximum likelihood in IQ-Tree v.3 (Wong et al. 2026); the ultrafast bootstrap was used to measure node confidence (Hoang et al. 2018). ModelFinder (Kalyaanamoorthy et al. 2017) was used to choose the optimum model for each tree based on Bayesian Information Criterion; the HKY+F+G4 model was chosen for cox1, and the TIM3+F+R3 model was chosen for the ribosomal tree. Figures were produced by exporting IQ-Tree files to the Interactive Tree of Life webserver (Letunic and Bork 2019).

### Population genetic inference

A 638 bp region of cox1 was sequenced in 33 new samples: 20 from the *H. heliophila* type location (10 each from the Rachel Carson Reserve and the Duke Aquafarm, both in Beaufort Harbor, North Carolina), seven from California, one from Virginia, three from Brazil, and two from France (Table S1). These samples were combined with publicly available data mined from GenBank and BOLD. Two previously sequenced samples were excluded due to apparent sequencing errors. One sample from Brazil (EU076812) and one from Argentina (LR880963) were excluded due to high numbers of apparent sequencing errors: both contain frame-shifting insertion/deletion mutations in the coding region together with many putative base changes that are not seen in any other sequence. To generate a haplotype network, sequences with ambiguous bases (bases called as N) and short sequences were removed from the alignment, and long sequences were trimmed in order to generate a 564 bp alignment with no missing data. This dataset included 174 sequences that were, to the best of our knowledge, all from unique samples. Heterozygosity (π) was calculated from this dataset by determining the mean proportion of pairwise differences between all DNA sequences in a sample. To determine the statistical significance of differences in π between samples, we used a permutation test as described below. The haplotype network was constructed with the haploNet function in the R package *pegas* (Paradis 2010).

## Results & Discussion

### Sponges matching the description of *H. heliophila* remain common at the type location

Wilson described the species *Stylotella heliophila* from material collected at Beaufort Harbor, North Carolina (*Stylotella* was later synonymized with *Hymeniacidon*). He stated, “This *Stylotella* is the most abundant sponge in Beaufort Harbor. Common on the bottom in shallow water attached to shells, also under wharves attached to piles, stones, etc.” The sponges were orange, sometimes with a greenish cast, and varied from encrusting forms to forms with erect lobes, with oscula at the ends of the lobes or on conical outgrowths. The surface had a roughened appearance, with small conules and visible subectosomal canals. The only spicules were styles, 120–350 × 4–8 μm. In the interior, these occurred both as scattered individual spicules and in reticulating multispicular tracts; spongin was not observed. The ectosomal skeleton contained styles that were “about horizontal” but projected slightly to form vague tufts and were sometimes arranged in loose tracts.

Sponges matching this description remain abundant throughout the Beaufort Harbor estuary. The Duke Aquafarm, an oyster-farming research and education facility located within the estuary, grows oysters in floating bags. These floating bags are commonly fouled by orange sponges (Figure 1B), which also encrust oysters on the tidal mudflats at this location (Figure 1A). The sponges have a roughened, conulose surface and range from thin encrustations to more upright growth forms, with oscula scattered across the surface or located at the tips of upright fistula. A second collection site within the estuary, the Rachel Carson Reserve, contains intertidal mudflats with extensive oyster beds (Figure 1C). As at the aquafarm, these oysters are commonly fouled by sponges matching Wilson’s description. We collected 12 specimens from each location and designated one specimen (TLT1736) as a topotype of *H. heliophila*. It has been deposited in the permanent collection TBD (will deposit after peer review).

**Figure 1.**
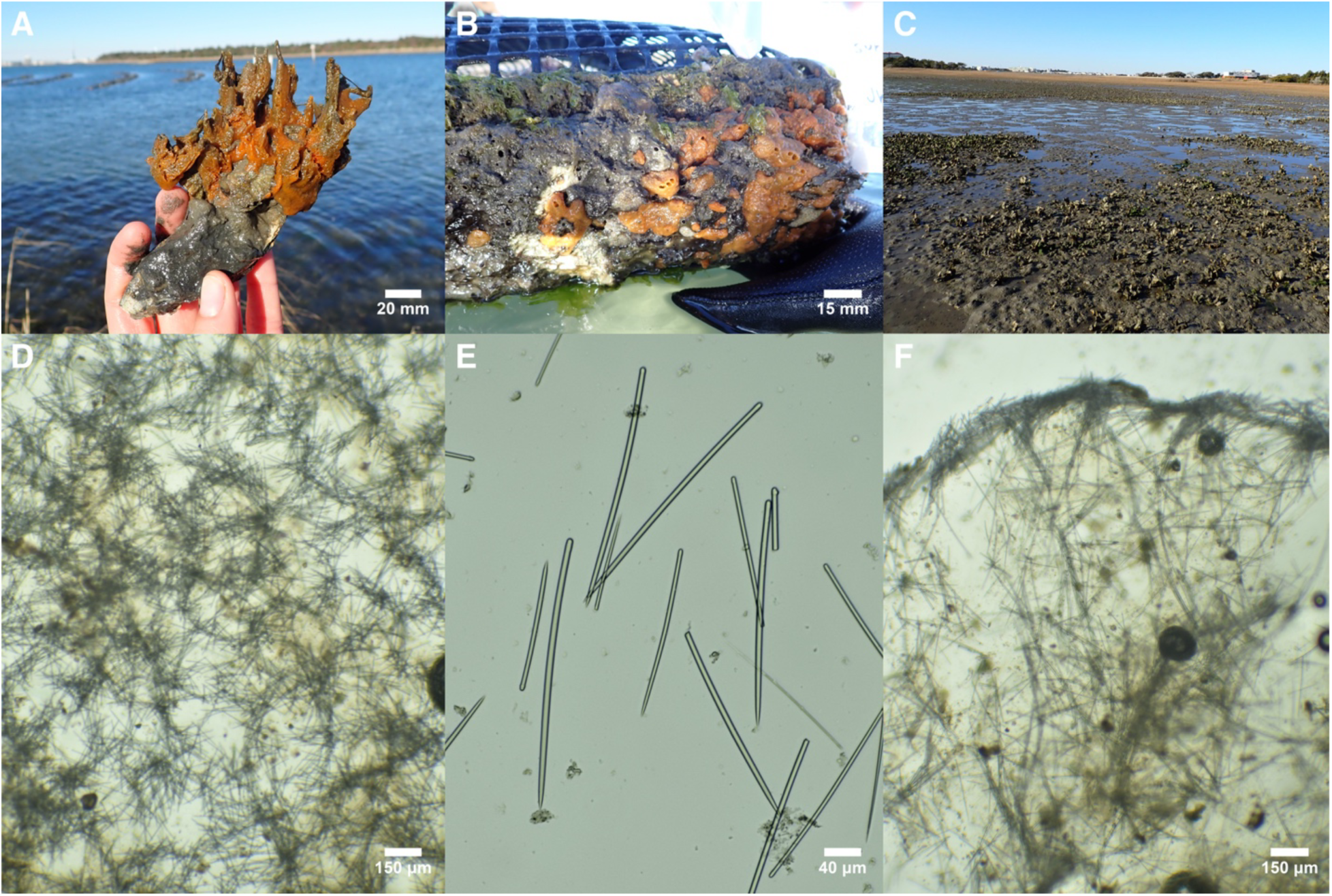
*Hymeniacidon* from Beaufort Harbor. A: Field photo of the *H. heliophila* topotype, TLT1736, growing on an oyster from an intertidal mudflat. B: Field photo of sample TLT1730 growing on a floating oyster bag. C: Site photo of the oyster reef at the Rachel Carson Reserve, where samples were collected from oysters exposed at low tide. D: Tangential section of the ectosome showing paratangential styles in bouquets. E: Styles, showing variation in length and tylote endings. F: Perpendicular section at sponge surface showing paratangential surface crust of styles, with multispicular tracts and styles in confusion in the choanosome. D-F are from the *H. heliophila* topotype, TLT1736.

The spicules of the topotype are exclusively styles (Figure 1E), varying considerably in both length and width, 114–238 × 2–7 μm (Table 1). The ectosomal skeleton contains loose paratangential bouquets of styles (Figure 1D), whereas the choanosome contains both individual styles in confusion and meandering multispicular tracts lacking visible spongin (Figure 1F). These features closely match Wilson’s original description. The styles range from straight and unmodified to forms with subtylote ends, with the tylote swelling sometimes displaced slightly from the spicule terminus toward the shaft (Figure 1E).

**Table 1.**
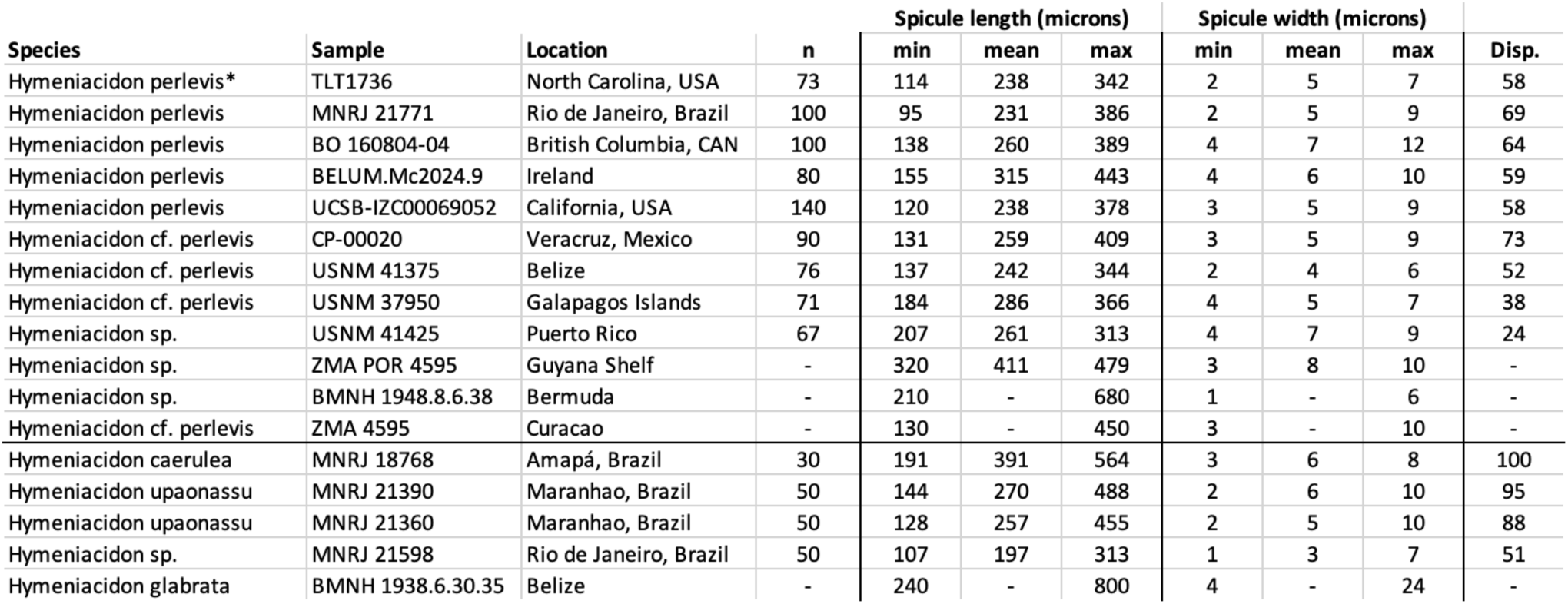
Spicule morphology. The asterisk indicates the sample that we designated as the *H. heliophila* topotype, which we subsequently synonymize with *H. perlevis*. Samples with genetic confirmation of species identify are listed as *H. perlevis*; samples with similar morphology but no genetic data are listed as *H.* cf. *perlevis,* and samples suspected to be other species are listed as *H. sp.* Disp. = root mean square dispersion. Samples with some values missing are taken from published papers.

| Species | Sample | Location | n | Spicule length (microns) |  |  | Spicule width (microns) |  |  | Disp. |
| --- | --- | --- | --- | --- | --- | --- | --- | --- | --- | --- |
|  |  |  |  | min | mean | max | min | mean | max |  |
| *Hymeniacidon perlevis* * | TLT1736 | North Carolina, USA | 73 | 114 | 238 | 342 | 2 | 5 | 7 | 58 |
| *Hymeniacidon perlevis* | MNRJ 21771 | Rio de Janeiro, Brazil | 100 | 95 | 231 | 386 | 2 | 5 | 9 | 69 |
| *Hymeniacidon perlevis* | BO 160804-04 | British Columbia, CAN | 100 | 138 | 260 | 389 | 4 | 7 | 12 | 64 |
| *Hymeniacidon perlevis* | BELUM.Mc2024.9 | Ireland | 80 | 155 | 315 | 443 | 4 | 6 | 10 | 59 |
| *Hymeniacidon perlevis* | UCSB-IZC00069052 | California, USA | 140 | 120 | 238 | 378 | 3 | 5 | 9 | 58 |
| *Hymeniacidon cf. perlevis* | CP-00020 | Veracruz, Mexico | 90 | 131 | 259 | 409 | 3 | 5 | 9 | 73 |
| *Hymeniacidon cf. perlevis* | USNM 41375 | Belize | 76 | 137 | 242 | 344 | 2 | 4 | 6 | 52 |
| *Hymeniacidon cf. perlevis* | USNM 37950 | Galapagos Islands | 71 | 184 | 286 | 366 | 4 | 5 | 7 | 38 |
| *Hymeniacidon sp.* | USNM 41425 | Puerto Rico | 67 | 207 | 261 | 313 | 4 | 7 | 9 | 24 |
| *Hymeniacidon sp.* | ZMA POR 4595 | Guyana Shelf | - | 320 | 411 | 479 | 3 | 8 | 10 | - |
| *Hymeniacidon sp.* | BMNH 1948.8.6.38 | Bermuda | - | 210 | - | 680 | 1 | - | 6 | - |
| *Hymeniacidon cf. perlevis* | ZMA 4595 | Curacao | - | 130 | - | 450 | 3 | - | 10 | - |
| *Hymeniacidon caerulea* | MNRJ 18768 | Amapá, Brazil | 30 | 191 | 391 | 564 | 3 | 6 | 8 | 100 |
| *Hymeniacidon upaonassu* | MNRJ 21390 | Maranhao, Brazil | 50 | 144 | 270 | 488 | 2 | 6 | 10 | 95 |
| *Hymeniacidon upaonassu* | MNRJ 21360 | Maranhao, Brazil | 50 | 128 | 257 | 455 | 2 | 5 | 10 | 88 |
| *Hymeniacidon sp.* | MNRJ 21598 | Rio de Janeiro, Brazil | 50 | 107 | 197 | 313 | 1 | 3 | 7 | 51 |
| *Hymeniacidon glabrata* | BMNH 1938.6.30.35 | Belize | - | 240 | - | 800 | 4 | - | 24 | - |

### Hymeniacidon heliophila is a junior synonym of Hymeniacidon perlevis

None of the morphological features of the *H. heliophila* topotype distinguish it from *H. perlevis*. The spicules are indistinguishable from those of *H. perlevis* specimens from distant regions (Table 1), and the skeletal architecture and other morphological features also match those of *H. perlevis* (Erpenbeck and van Soest 2002; Turner 2020; Turner et al. 2025). These observations are consistent with *H. heliophila* being a synonym of *H. perlevis*. Also consistent with these data is the possibility that the two taxa are cryptic species that are morphologically similar but evolutionarily distinct. To distinguish between these possibilities, we sequenced the Folmer barcoding region of the cox1 locus from 10 specimens collected at each of the two Beaufort Harbor sites. All 20 sequences were identical across the entire 638-bp region analyzed, providing no evidence for the presence of multiple species within the estuary (Figure 2). Furthermore, these sequences are identical to the most common worldwide haplotype of *H. perlevis* (Figure 3).

**Figure 2.**
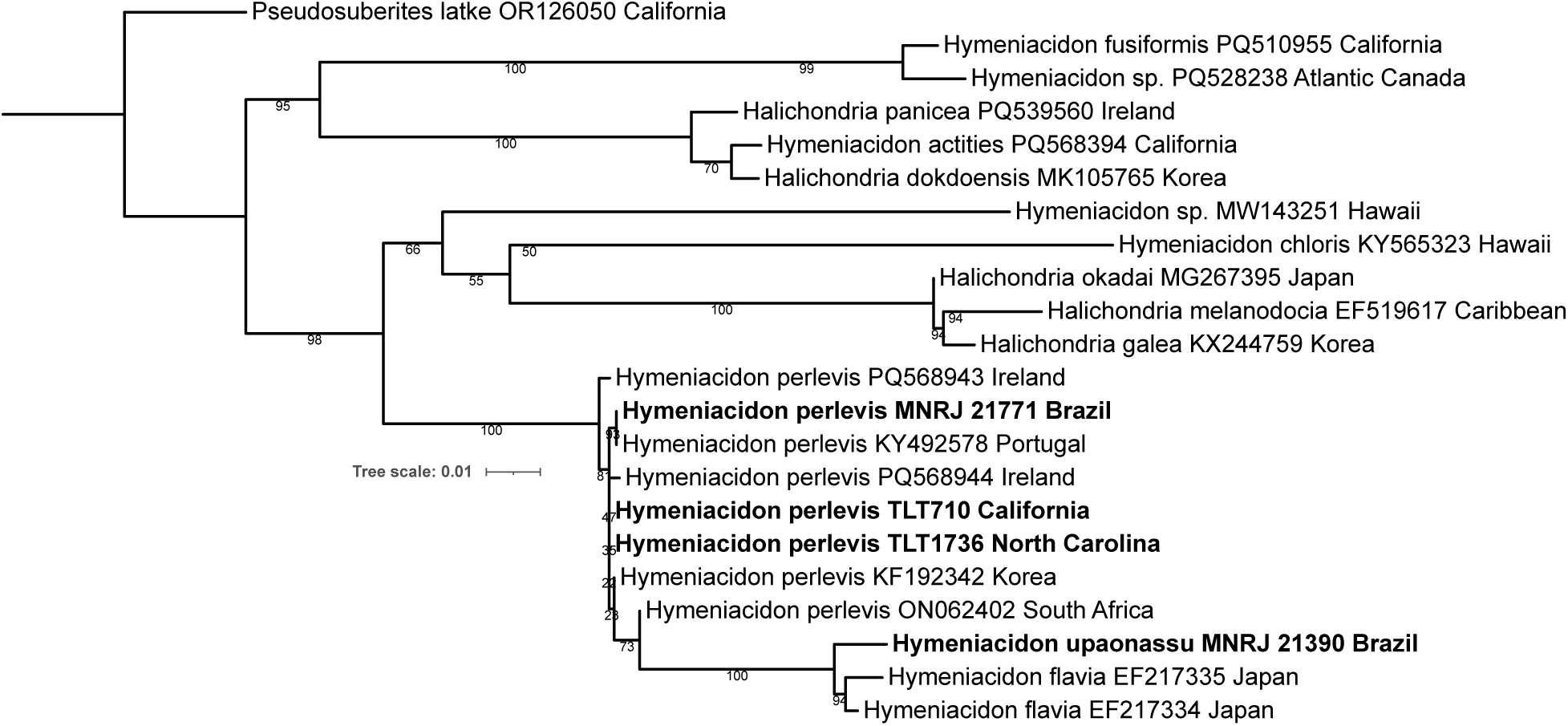
Maximum likelihood phylogeny of the cox1 locus. Bold indicates new sequences; because of the large number of identical sequences, only select representatives of *H. perlevis* are shown, along with other *Hymeniacidon* species and outgroups. Sample names are included for new sequences, and GenBank accession numbers are included for previously published sequences. Node confidence is based on bootstrapping. Scale bar indicates substitutions per site. Sample TLT1736 is the *H. heliophila* topotype, subsequently synonymized with *H. perlevis*.

**Figure 3.**
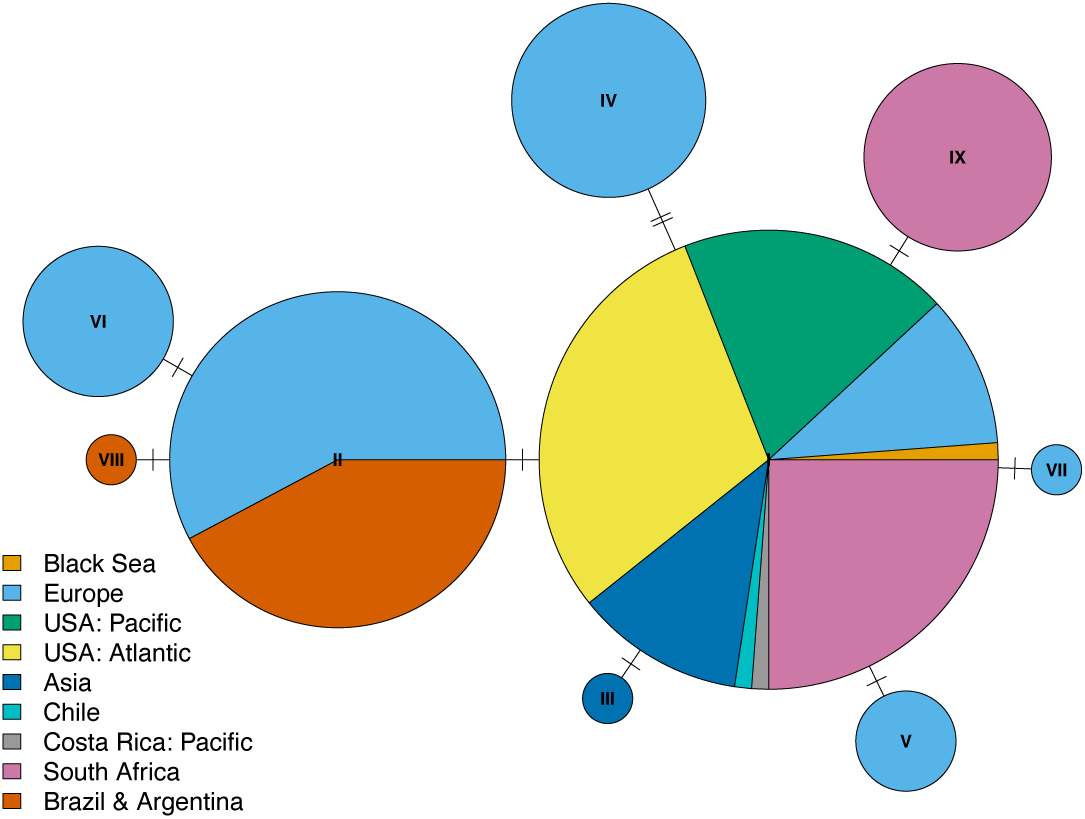
Haplotype network at the cox1 locus. Each circle is a unique haplotype, with the size of the circle proportional to the number of samples; genotypes III, VII, and VIII are singletons. The tick marks linking haplotypes indicate the number of genetic differences between them. The network includes 174 samples aligned across a 564 bp region, including 34 newly generated sequences. The 20 sequences from the *H. heliophila* type location (Beaufort Harbor, North Carolina) are included in the “USA: Atlantic” portion.

The cox1 locus is known to evolve slowly in sponges (Huang et al. 2008; López-Legentil et al. 2010; Pöppe et al. 2010; Turner and Pankey 2023). To further test whether the common *Hymeniacidon* in Beaufort Harbor is *H. perlevis*, we also sequenced portions of the nuclear ribosomal locus. For the topotype and three additional specimens, we sequenced three adjacent amplicons spanning 2,304 bp, including the most rapidly evolving regions of the locus (the internal transcribed spacer and the 28S C1-D1 barcoding region). We also sequenced subsets of this region from specimens collected in California, Virginia, Brazil, and Europe. The four Beaufort Harbor specimens, including the topotype, were genetically identical. As shown in Figure 4, the ribosomal phylogeny places these specimens firmly within a clade of *H. perlevis* from around the world.

**Figure 4.**
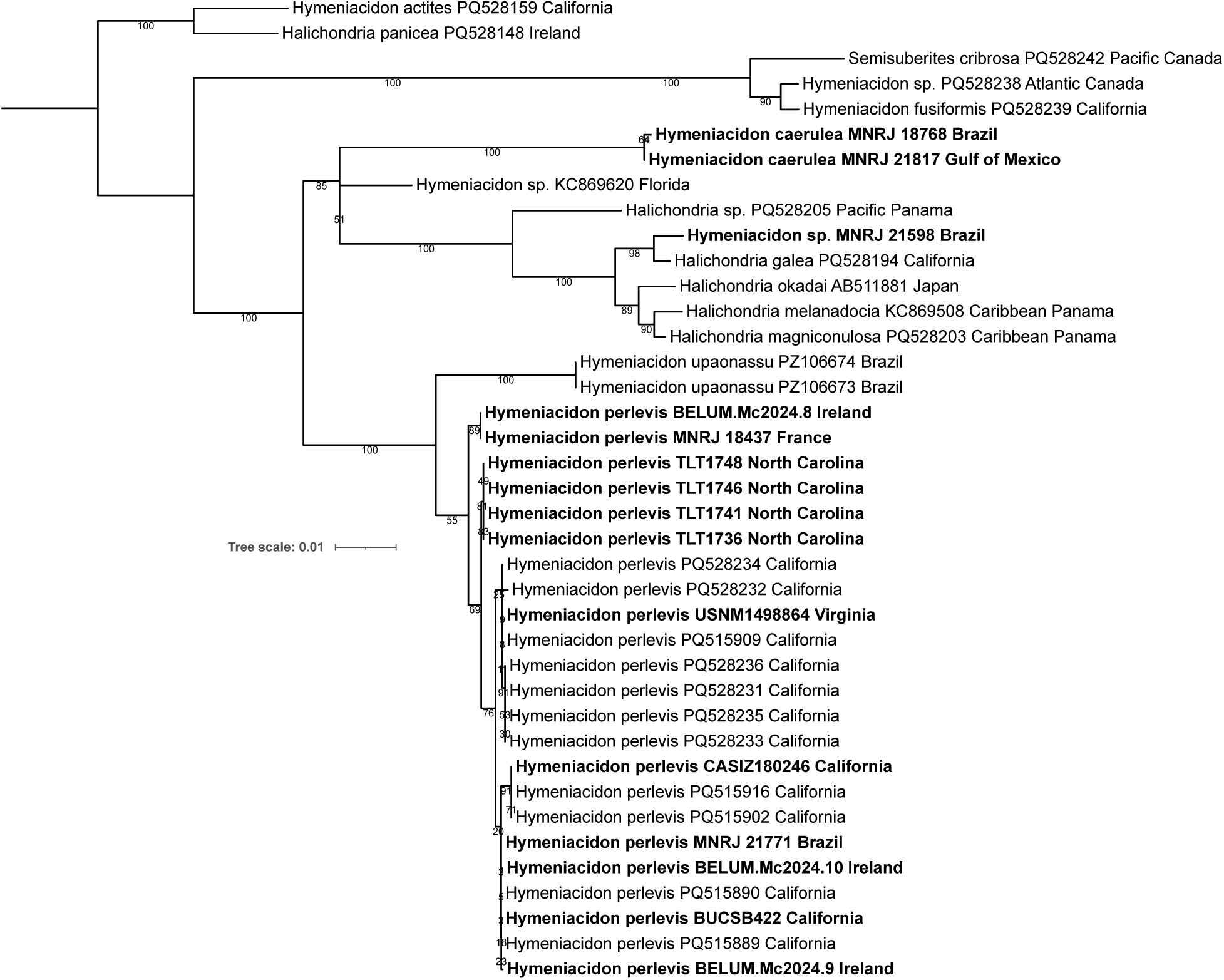
Maximum likelihood phylogeny of the nuclear ribosomal locus. Bold indicates new sequences. Sample names are included for new sequences, and GenBank accession numbers are included for previously published sequences. Node confidence is based on bootstrapping. Scale bar indicates substitutions per site. Sample TLT1736 is the *H. heliophila* topotype, subsequently synonymized with *H. perlevis*.

Together, these morphological and genetic data strongly support the conspecificity of the common *Hymeniacidon* in Beaufort Harbor and *H. perlevis*. It remains possible that the sponge examined by Wilson in 1911 belonged to a different species that was morphologically indistinguishable from *H. perlevis* and has since been replaced by an introduced population of *H. perlevis*. We consider this scenario unlikely, and it cannot be tested because no material from Wilson’s collections is available for genetic analysis. We therefore recommend treating *H. heliophila* as a junior synonym of *H. perlevis*.

### Hymeniacidon perlevis likely originated in Europe

It is highly unlikely that *H. perlevis* achieved its remarkably broad global distribution without human-mediated transport. Because population bottlenecks associated with species introductions are expected to reduce genetic diversity, we used the cox1 dataset assembled here to investigate the species’ native range. We divided the available samples into six broad geographic regions (Table 2). Genetic diversity in Europe (π = 0.29%) was substantially higher than in any other region examined (0.00–0.09%). To evaluate the statistical significance of this result, we performed a permutation test. The locations of all 174 samples were randomly reassigned 1,000 times, and a subset of 64 samples (equal to the European sample size) was drawn from each randomized dataset. Genetic diversity in the permuted datasets ranged from 0.12–0.25%, and none equaled or exceeded the observed European value of 0.29%, indicating a highly significant difference (p < 0.001).

**Table 2.** Nucleotide diversity (π) at cox1. SNPs = single nucleotide polymorphisms. “World excluding Europe” includes three samples not included in any of the sub-regions (one each from Pacific Costa Rica, Chile, and the Black Sea).

|  | N | All variability |  | Excluding singletons |  |
| --- | --- | --- | --- | --- | --- |
| | | $\pi$ | SNPs | $\pi$ | SNPs |
| Europe: Portugal, France, Ireland, Netherlands | 64 | 0.29% | 5 | 0.29% | 5 |
| South Africa | 35 | 0.09% | 1 | 0.09% | 1 |
| Asia: Korea, China, Japan | 11 | 0.03% | 1 | 0.00% | 0 |
| SW Atlantic: Brazil, Argentina | 20 | 0.03% | 2 | 0.00% | 0 |
| US Pacific: California | 16 | 0.00% | 0 | 0.00% | 0 |
| US Atlantic: North Carolina, Virginia, Florida | 25 | 0.00% | 0 | 0.00% | 0 |
| World excluding Europe | 110 | 0.10% | 4 | 0.10% | 2 |

The contrast between Europe and the rest of the world becomes even stronger when singleton variants are excluded. Singletons are single nucleotide polymorphisms (SNPs) observed in only one of the 174 samples. Although some singletons likely represent genuine genetic variation, sequencing errors are expected to be rare and randomly distributed and therefore will disproportionately appear as singleton variants. Excluding singletons has no effect on European diversity because all five variable sites occur in multiple individuals. In contrast, only a single variable site remains in any region outside Europe: a common SNP in South Africa that is private to that population.

These results support a European origin for *H. perlevis* and suggest that all sampled populations outside Europe are introduced. The data may also indicate an independent introduction to Brazil and Argentina. A SNP that is polymorphic in Europe is fixed for one allele in Brazil and Argentina but fixed for the alternative allele in all other sampled regions (Figure 3). Although multi-locus genetic data would strengthen these conclusions, no other locus currently has comparable geographic sampling both within and outside Europe.

### Hymeniacidon perlevis remains unconfirmed in the Caribbean

Having established that *H. heliophila* is a synonym of *H. perlevis*, we next investigated the extent to which *H. perlevis* extends into the warm waters in the central western Atlantic. DNA data generated here confirms the species is present in Virginia and North Carolina; when combined with previously sequenced records from Palm Beach, Florida (Erpenbeck et al. 2007), it becomes clear that the species has a considerable range on the Atlantic coast of the Southeastern USA (Figure 5). Genetic data also confirms the species at a single location in the northern Gulf of Mexico: Dauphin Island, Alabama (Erwin et al. 2011). We do not expect this to be an isolated population, as images posted to the site iNaturalist.org show that sponges consistent with the morphology of *H. perlevis* are found from throughout the gulf coast of Florida at least as far south as Fort Myers.

**Figure 5.**
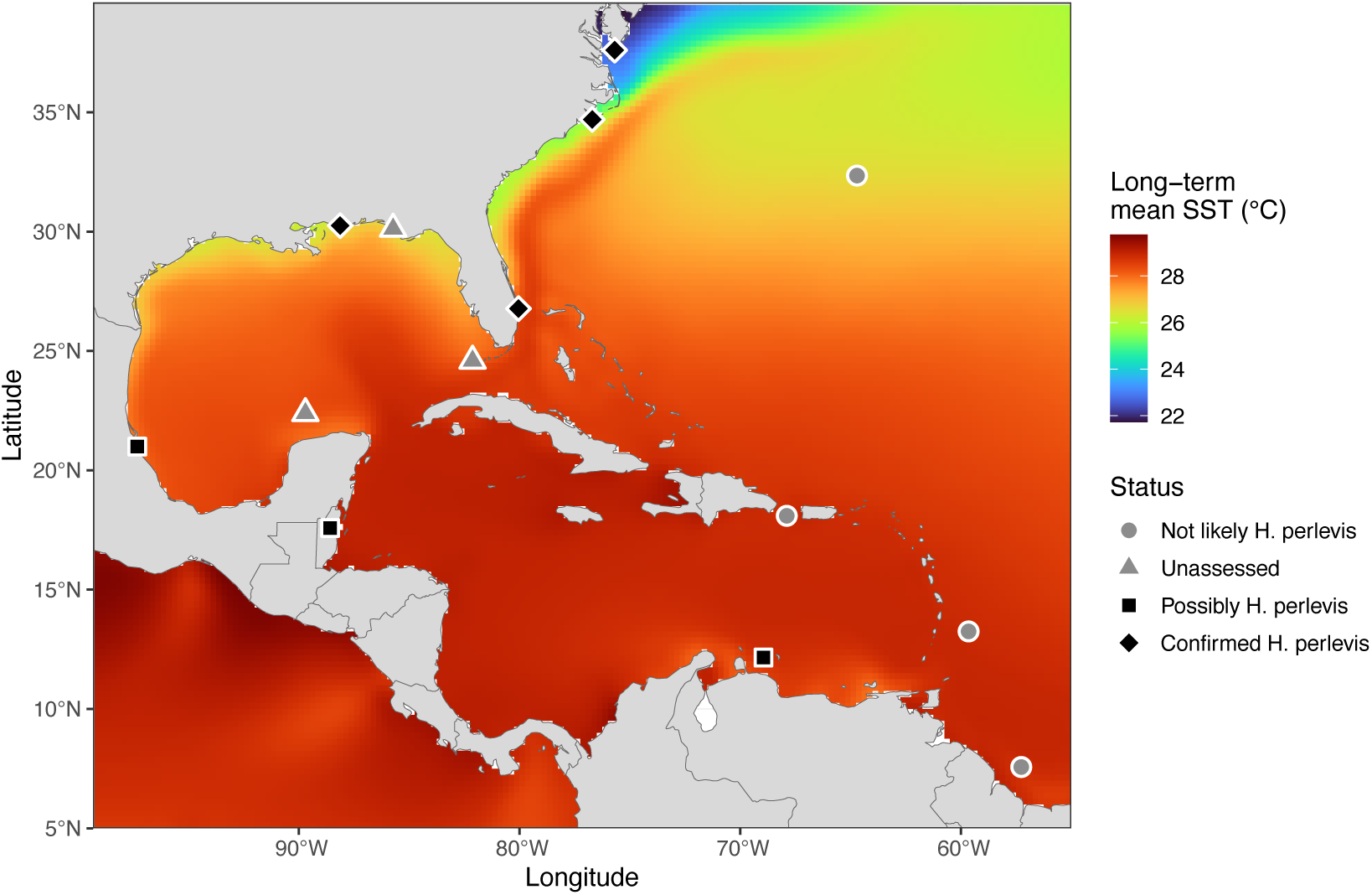
*Hymeniacidon* samples from the central western Atlantic. All samples genetically confirmed as *H. perlevis* are shown as black diamonds. Circles and squares are samples that were examined, or assessed from published data, and deemed to be morphologically consistent with *H. perlevis* (squares) or not (circles). Triangles are samples mentioned in the text and not assessed here, but worthy of future examination. Colors indicate long-term sea surface temperature from the NOAA OISST v2 high-resolution monthly long-term-mean (1991-2020 climatology).

In contrast to these confirmed reports in Florida and the Gulf of Mexico, there is scant evidence of the species in the Caribbean. We know of no samples with genetic confirmation south of West Palm Beach, Florida until reaching the state of Rio de Janeiro, Brazil (Figures 3 & 4). We attempted to confirm the species in the Caribbean by acquiring previously vouchered *Hymeniacidon* samples from the Smithsonian Institution that were collected in Puerto Rico, Belize, and Barbados (Tables 1, S1). We attempted to sequence the cox1 and 28S loci from these samples, but like many older sponge samples, they did not appear to have well-preserved DNA, and we were not successful.

With genetic data unavailable, we investigated the morphology of samples from the Caribbean and surrounding regions. Several morphological features were considered. Field photos are useful, despite the high variability in growth form, because some other species in the region are distinguished by color: *H. caerulea* is blue and *H. upaonassu* is light-yellow (Fortunato et al. 2020). All known cases of *H. perlevis* in the region are orange (Lôbo-Hajdu et al. 2004; Muricy and Hajdu 2006; Ribeiro et al. 2010), though this species is known to be yellow, pink, or red in other regions of the world (Erpenbeck and van Soest 2002; Ackers et al. 2007; Samaai et al. 2022; Turner et al. 2025). Spicule measurements are also useful in diagnosing the species (Table 1). In addition to mean values, the range in spicule sizes is sometimes able to differentiate among species of *Hymeniacidon* (Turner et al. 2025). *H. perlevis* has a large range in spicule sizes due, in part, to having smaller spicules in the ectosome vs. the choanosome. Some reports consider the species to have two or three size-classes of spicules (Gastaldi et al. 2018; Samaai et al. 2022), but samples that we have examined have a continuous distribution of both length and width (Figure 6). We do find that the spicules are significantly shorter in the ectosome, but the full range of lengths is present in both the ectosome and choanosome, so dividing spicules into classes requires arbitrary partitions. As an alternative approach, we calculated the dispersion in these values in two-dimensional space as the root mean square Euclidean distance of individual spicules from the specimen centroid (see methods). Only summary statistics are usually reported in the sponge taxonomy literature, but authors of two past reports (Harbo et al. 2021; de la Cruz-Francisco 2025) shared their raw data with us, which allowed for this calculation to be added to Table 1.

**Figure 6.**
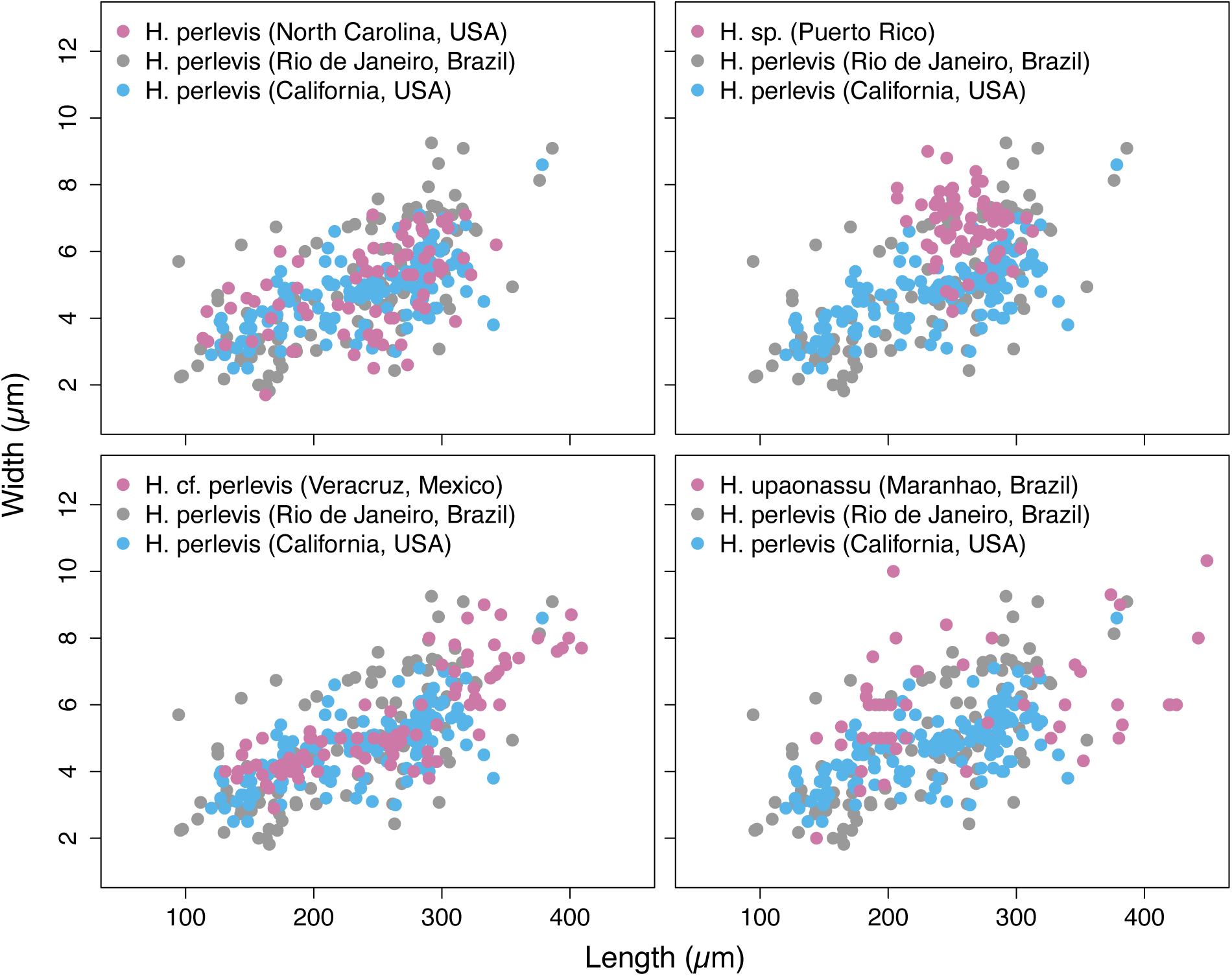
Spicule variation in *Hymeniacidon*. Upper left: Samples of *H. perlevis* have wide variation in spicule length and width. A sample from North Carolina (the *H. helophila* topotype, TLT1736, in pink) is extremely similar samples from California (UCSB-IZC00069052) and Brazil (MNRJ 21771). The other panels compare *H. perlevis* spicules to samples without genetic data from Puerto Rico (USNM 41425, unlikely to be *H. perlevis* due to low dispersion) and Mexico (CP-00020, likely *H. perlevis*), as well as the sympatric species *H. upaonassu* (MNRJ 21390, Brazil).

Using these morphological features, we divided samples without genetic data into those that were consistent with the morphology of *H. perlevis* and those that were likely other species. The sample examined from Barbados (USNM 42448) contained an additional type of spicule (tylotes with dimpled ends), so this sample was not in the genus *Hymeniacidon*. The sample from Puerto Rico had styles with very similar mean values as *H. perlevis*, but the dispersion of these values was much lower (Figure 6), so this sample is likely to be a different species of *Hymeniacidon*. The spicules of this sample do not match any of the other *Hymeniacidon* species from the region (*H. glabrata*, *H. upaonassu,* and *H. caerulea*, Table 1), so it is likely an undescribed species. We also consider past reports of *H. heliophila* from the Guyana Shelf (van Soest 2017) and Bermuda (de Laubenfels 1950) to be unlikely to be *H. perlevis*. Though we did not reexamine these samples ourselves, the previously reported mean and/or maximum spicule lengths, and the lack of short spicules, seem likely to make these samples members of a different species (Table 1). In contrast, samples reported from Tuxpan, Mexico, on the Western shores of the Gulf of Mexico, are morphologically consistent with *H. perlevis* (de la Cruz-Francisco 2025). Field photos of living sponges match the color and other morphological features of *H. perlevis*, the reported spicule dimensions and dispersion of these values are consistent with *H. perlevis* (Table 1, Figure 6). An additional sample from Belize (USNM 41375) has less data available, but could be *H. perlevis*: though this sample lacks field photos, the mean and dispersion of spicule sizes is consistent with this species. A previous report of *H. perlevis* from Curaçao (Table 1) lacks field photos, dispersion data, and even mean values, but the spicule ranges reported are at least consistent with *H. perlevis*, so we also consider this is a possible, if extremely preliminary, record of the species. (Though outside the scope of this investigation, we also examined a sample from the Galapagos Islands, Ecuador, and these data are included in Table 1.)

Together, these data illustrate that *H. perlevis* is likely to be broadly distributed in the coastal Gulf of Mexico. Additional vouchers that were not examined here may extend this range slightly to the South: The University of Florida Museum has vouchers (e.g. UF 2435) identified as *H. perlevis* from the Florida Keys, and the Colección Nacional del Phylum Porifera “Gerardo Green”, located at the Universidad Nacional Autónoma de México, has a sample identified as *H. perlevis* from the Yucatan peninsula (CNPGG 1412). In contrast, few records are available from the Caribbean, and many of those appear unlikely to be specimens of *H. perlevis*. Those that are consistent with the morphology of *H. perlevis* could be different species with similar morphology. Supporting this possibility, one sample we examined from Rio de Janeiro appeared consistent with the morphology of *H. perlevis*, but the 28S sequence places the sample in a distant part of the phylogeny (MNRJ 21598, Figure 4, Table 1). This sample appears more closely related to *Halichondria galea* than to any other species of *Hymeniacidon*. This result is surprising, and performing a replicate DNA extraction on this sample to rule out the possibility of DNA contamination would be desirable as confirmation. The result is not implausible, however, as *Hymeniacidon actites* was recently found to be closely related to *Halichondria dokdoensis*, showing that recent evolution of oxeas into styles has taken place (Turner et al. 2025; Figures 3 & 4).

## Conclusions

For over a century, *Hymeniacidon heliophila* has been used to study symbiosis, regeneration, ecology, and evolution in sponges (Parker 1910; de Laubenfels 1947; Turque et al. 2008; Ribeiro et al. 2010; Weigel and Erwin 2016; Coutinho et al. 2017; Ascer et al. 2025). This body of research can now be united with studies conducted on *H. perlevis* from Europe and other places, as these populations are members of the same species, and should be hereafter referred to as *H. perlevis*. This species was first described in Europe (Montagu 1814), and population genetic analysis of the cox1 locus suggests that Europe includes the native range of the species, with all other genotyped populations the result of human introductions. These introductions are not especially recent, with the sponge already abundant in a North Carolina estuary by 1910 (Parker 1910) and abundant in a Southern California estuary by 1914 (Harbo et al. 2021). The strength of these conclusions is limited by having only a single mitochondrial marker with comparable data, and obtaining comparable multi-locus data should be a high priority for future work.

The ecological breadth of *H. perlevis* is impressive in multiple ways, and the dramatic range of temperatures in which it thrives is one notable example. However, our results make it seem unlikely that the species is present in the Caribbean. We are aware of few reports of the species from the region, and closer examination of some of these vouchered specimens reveal that they are likely to be members of other species (Figure 5). Only a few samples from the region have morphological data potentially consistent with the species, and it is plausible that these are members of morphologically similar species that await description.

The global distribution of *H. perlevis*, together with its abundance in places that are easy to sample, make it among the most promising species to use as a model system for marine sponges. We hope the results presented here motivate further work on the ecology, evolution, and applied potential of the species. We also hope that recognition of its introduced status will motivate work investigating its potential impact on the ecological and economical services of the affected ecosystems.

## Supporting information

Table S1. Metadata for all samples investigated.

Table S2. Measurement data from all spicules.

## Acknowledgments

We thank Abigail Reft and Allen Collins for loaning samples from the Smithsonian National Museum of Natural History collection. California collections were facilitated by many members of the UCSB’s Marine Science Institute, especially Skylah Reis, Robert Miller, Clint Nelson, Christoph Pierre, and Christian Orsini. Collection and photography of North Carolina samples were facilitated by Joshua Osterberg and Emma Johnson. We thank Julio C. C. Fernandez for spicule photographs of MNRJ samples and Thiago S. de Paula for molecular preparation and sequencing of MNRJ samples. We thank Thomas Schultz for management of the Duke Aquafarm.

## Statements and Declarations

### Author Contributions

TLT, HFMF, and GLH conceived and designed the experiments, and TLT, HFMF, and JMW collected samples and data. TLT performed the analyses. All authors were involved in producing the manuscript.

## Supplementary Data

Table S1. Metadata for all samples investigated. Species = species name. Authority = Citation of paper describing species. Type status = Holotype, Paratype, or neither (NA). Sample ID: a unique identifier used for newly collected samples; samples with a museum voucher but no collection ID were loaned by the corresponding museum without information regarding the sample ID. Date = collection date. Collection location = text description of collection location. Collection region = state/province of collection location. Lat and Long: Latitude and Longitude of collection location, when known. Depth = collection depth, when known. Mitochondrial voucher = GenBank ID for sequenced mitochondrial DNA; NA indicated no DNA was sequenced. Ribosomal voucher = GenBank ID for sequenced nuclear ribosomal DNA. Collector = collector, when known. Notes = additional information.

**Table S2. Measurement data from all spicules. Lengths and widths are provided for each spicule, in microns. Sample ID or voucher number = sample ID (TLT number) for newly collected samples, or museum voucher number for loaned vouchers.**

